# Nucleation Landscape of Biomolecular Condensates in the Grand Canonical Ensemble via Monte Carlo Simulations

**DOI:** 10.1101/2025.06.06.658304

**Authors:** Aliasghar Sepehri, Gül H. Zerze

**Affiliations:** Department of Chemical and Biomolecular Engineering, University of Houston, Houston, Texas, 77204, USA

## Abstract

We present a Monte Carlo framework for simulating the nucleation of biomolecular condensates in the grand canonical ensemble, which overcomes the limitations of fixed particle number and allows direct control of the dilute-phase concentration. Our approach combines conformation sampling with bias-enhanced cluster size sampling, enabling accurate sampling of individual cluster sizes in various conformations. Our method resolves nucleation free energy surfaces and is capable of capturing both classical and nonclassical nucleation mechanisms. We validated our method by reproducing structural properties of disordered proteins across a diverse benchmark set and by applying it to study nucleation in two phase-separating proteins, FUS-LC and NDDX4, using both HPS and MPIPI coarse-grained force fields. While both proteins exhibit classical nucleation behavior under the HPS model, only FUS-LC remains classical with MPIPI. In contrast, NDDX4 shows a distinctly nonclassical nucleation pathway under MPIPI, characterized by a metastable intermediate state near the cluster size of 53 protein chains. Morphological analysis reveals that clusters up to this point are compact and spherical, whereas larger clusters adopt a nonspherical, two-lobed geometry indicative of frustrated coalescence. These findings underscore the critical role of sequence composition and force field parameterization in shaping nucleation pathways and demonstrate the utility of our framework for uncovering complex, mechanism-rich free energy landscapes in biomolecular condensation.

## Introduction

Biomolecular condensates are membraneless organelles that serve as dynamic compartments, facilitating essential biological processes such as DNA replication, DNA repair, and transcription by concentrating specific biomolecules, including proteins, RNA, and DNA.^1,2^ These condensates are formed primarily by liquid-liquid phase separation (LLPS), leading to the emergence of distinct dilute and condensed phases.^2–5^ These assemblies are typically driven by weak multivalent interactions that involve intrinsically disordered regions (IDRs) and folded domains.^2,6,7^ Although the thermodynamics of phase separation have been increasingly mapped out,^8–10^ our understanding of the kinetics and nucleation mechanisms of condensate formation remains incomplete.

In metastable regimes, condensates form via nucleation, a process that requires overcoming a free energy barrier to create a critical-sized cluster. This stands in contrast to spinodal decomposition, which proceeds spontaneously in systems beyond the limits of metastability.^11–13^ Molecular simulations of nucleation present two significant challenges, both of which can be effectively addressed using grand canonical Monte Carlo (GCMC) simulations. The first challenge is finite-size effects in the canonical ensemble, which results in alteration of the solution phase density and the chemical potential due to depletion during nucleation, although nucleation theories, such as the classical nucleation theory (CNT), assume an infinite reservoir of particles in the supersaturated solution (or mixture), maintaining a constant density.^14^ To address this limitation, the modified liquid droplet (MLD) model was developed to account for particle depletion in the vapor phase in CNT.^15,16^ This approach has been successfully applied to biomolecular condensate nucleation simulations, too. ^17^ In our work, we employ the grand canonical ensemble (*µV T*), where the chemical potential (*µ*), corresponding to the initial phase density, is fixed, allowing particles to be added to or removed from the simulation system as needed.^18^

However, MLD remains tied to the assumptions of CNT and relies on fitting a predefined functional form to the free energy barrier. As such, it does not allow for direct, assumption-free exploration of the full nucleation landscape—particularly in systems where nonclassical behavior may emerge. To overcome this, we employ the grand canonical ensemble (*µV T*), where the chemical potential (*µ*), corresponding to the initial phase density, is fixed, allowing particles to be added to or removed from the simulation system as needed.^18^ This ensemble allows us to directly probe nucleation mechanisms without constraining the results to a classical form.

The second challenge is simulating nonclassical nucleation. Unlike classical nucleation, which involves a single critical nucleus, nonclassical nucleation proceeds through metastable intermediate states. This leads to complex free energy landscapes characterized by multiple maxima or saddle points, indicating alternative nucleation pathways.^19–22^ Some previous studies adopted models similar to CNT, aiming to determine physical parameters such as dilute density, condensed density, and surface tension.^17^ Our approach ensures comprehensive sampling of each cluster size, facilitating the identification of metastable intermediate clusters.

In the methodology section, we describe our approach, which integrates several Monte Carlo techniques, including aggregation-volume-bias Monte Carlo (AVBMC),^23–25^ configurational-bias Monte Carlo (CBMC),^26–29^ fixed-endpoints configurational-bias Monte Carlo (FECBMC),^30–35^ and umbrella sampling.^36^ AVBMC enhances the sampling of cluster formation by efficiently handling molecular aggregation, while CBMC and FECBMC improve conformational sampling by generating physically relevant molecular structures. Umbrella sampling is employed to overcome free energy barriers, ensuring accurate representation of rare events. This combined method has been shown to reproduce experimental results with high accuracy.^37^ Moreover, it can identify metastable states, such as the saddle point at a cluster size of five in the nucleation free energy profile of water, which arises due to stable hydrogen bonding interactions.^38^ Although this method has been primarily applied to vapor-liquid nucleation, it can also be used for nucleating condensates with implicit solvent force field models, as the implicit nature of the solvation makes the liquid-liquid phase separation of condensates analogous to vapor-liquid phase transitions, and other vapor-liquid equilibrium sampling techniques have been extensively used in studying the phase behavior of condensates. ^39–43^ This adaptability makes the approach valuable for studying a broader range of phase behavior phenomena, including biomolecular condensation and other self-assembly processes.

### Methodology

#### Overview

We developed a GCMC framework for mapping the nucleation free energy landscapes of biomolecular condensates. Our approach combines several advanced Monte Carlo techniques, including AVBMC, CBMC, and FECBMC, to enhance sampling efficiency across conformational space and cluster sizes. This section details the statistical mechanics basis and implementation of these techniques.

#### Free Energy Estimation of Clusters

If the formation of a cluster of size *n* is assumed to occur through a reversible aggregation of *n* monomers, while neglecting interactions between clusters, the formation Gibbs free energy difference associated with cluster formation is given by^14^

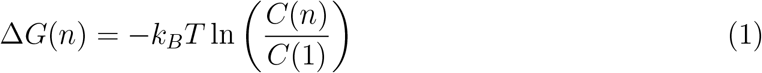

where *C*(*n*) and *C*(1) represent the equilibrium concentrations of clusters of size *n* and monomers, respectively. The free energy of a monomer is taken as the reference point and is therefore set to zero. This expression is valid in the regime where cluster sizes are smaller than the critical nucleus size *n*^*^, which corresponds to the maximum in the free energy profile.

Eq. 1 is particularly useful for simplifying umbrella sampling by reducing it to a single window that includes all possible cluster sizes. In this framework, the biasing potential at cluster size *n* is set to −Δ*G*(*n*).^36^

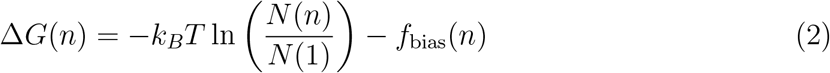

where *N* (*n*) and *N* (1) denote the number of times clusters of size *n* and monomers are sampled during the MC simulation, respectively. Initially, the biasing potential *f*_bias_(*n*) is set to zero for all cluster sizes. After each iteration of the simulation, the biasing potential is updated based on the cluster size histogram:

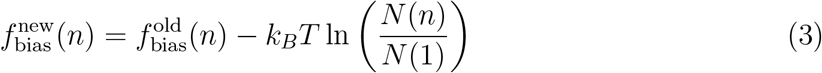

The simulation is considered converged when all cluster sizes are sampled uniformly. We also note that only a single cluster is simulated (using non-periodic boundary conditions), and any MC move that breaks this cluster is rejected. Since the simulation is conducted in the grand canonical ensemble, different cluster sizes can be sampled via particle insertions into or deletions from the cluster.

Since Eq. 1 is only valid for *n < n*^*^, the dilute density *ρ* and the maximum cluster size *n*_max_ must be chosen such that *n*_max_ *< n*^*^. Once convergence is achieved at a given dilute density *ρ*, the free energy profile at a different dilute density *ρ*^′^ can be calculated by^44^

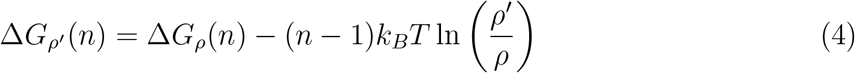

#### Cluster Definition

In previous simulations involving small molecules,^38^ cluster definitions were based on either distance cutoffs or interaction energy thresholds—i.e., two molecules were considered part of the same cluster if their interaction energy was below a specified cutoff, or if the distance between their centers of mass (or between their first atoms) was less than a defined threshold. However, since this study focuses nucleating dense phases of biomacromolecules, we employed a minimum atomic distance criterion instead: two molecules are considered part of the same cluster if at least one pair of beads—each from a different molecule—lies within a cutoff distance (e.g., 5 Å).^45^

#### Monte Carlo Moves

##### Translation and Rotation

We employed traditional translation and rotation moves to generate rigid-body configurations from a given state, enabling sampling of the relative positions and orientations of molecules within the cluster. In each MC step, a molecule is randomly selected and either translated by a random displacement or rotated by a random angle to produce a trial configuration. The probability of accepting this new configuration is evaluated using the Metropolis criterion: ^46^

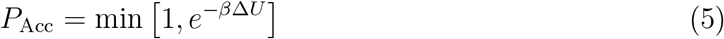

where Δ*U* is the change in potential energy between the new and old states, and *β* = 1*/*(*k*_*B*_*T*). Here, *k*_*B*_ is the Boltzmann constant and *T* is the absolute temperature of the system. The new configuration is accepted if *P*_Acc_ exceeds a uniformly generated random number between 0 and 1.

##### Configurational-Bias Monte Carlo (CBMC) and Fixed-End CBMC (FECBMC)

In the CBMC method,^26–28^ a randomly selected molecule is cut at a randomly selected segment and regrown, segment by segment, to reconstruct a new conformation. To preferentially generate conformations with lower energies, the Rosenbluth and Rosenbluth scheme, ^47^ originally developed to avoid overlaps in lattice models, is employed. During the regrowth of each segment, *k*_trial_ trial orientations are generated, and one is selected with a probability proportional to its Boltzmann factor. Both bonded (e.g., bond stretching, bond angle bending, torsion angle) and non-bonded (e.g., Lennard-Jones and electrostatic) interactions are considered. Since bonded interactions are significantly stronger and more restrictive, it is more efficient^29^ to generate trial orientations according to the Boltzmann distribution of the bonded energy *U* ^B^, while applying the Rosenbluth scheme only to the non-bonded energy *U* ^NB^. For each segment, *k*_trial_ orientations are generated from the distribution exp(−*βU* ^B^), and one is selected with the probability

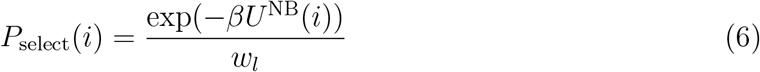

where

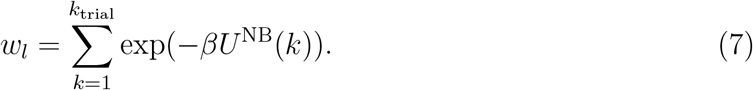

The overall Rosenbluth weight for growing *n*_seg_ segments is given by

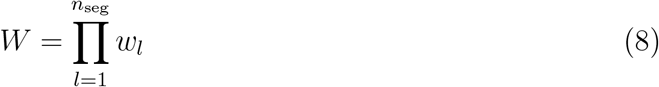

This regrowth procedure is repeated for the existing (old) conformation, with the difference that the first trial orientation corresponds to the current configuration of the segment. In this case, only *k*_trial_ − 1 new orientations are generated, and the first trial (the old configuration) is always selected. The acceptance probability for the new conformation is then given by

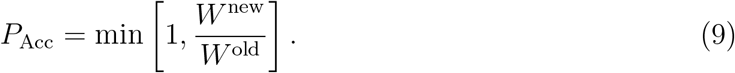

The acceptance probability decreases as the number of regrown segments increases (also see Fig. 5). Consequently, sampling the conformations of internal sections in long biomolecules becomes prohibitively less efficient, especially for larger polymers. To address this issue, we employed the FECBMC move. ^30–35^ In this method, two segments are randomly chosen as fixed points, the intermediate segments are deleted and then regrown to generate a new conformation. Since the biomolecular force fields used in this study lack bending and torsional potentials (as discussed in the next section), the algorithm developed by Escobedo and de Pablo^31^ is particularly efficient and suitable for our purposes. The acceptance probability for the new conformation is given by

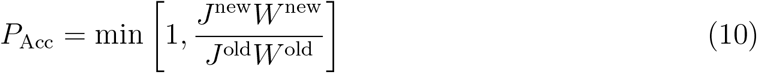

where *J* is the Jacobian factor that arises due to the geometric constraints imposed by regrowing the chain between two fixed points.

##### Aggregation Volume Bias Monte Carlo (AVBMC)

To sample a wide range of cluster sizes, swap moves are employed to insert or delete molecules. In the traditional swap move, insertion is performed by placing a molecule at a random position within the simulation box, while deletion involves randomly selecting and removing a molecule from the system.^48^ However, such randomly generated configurations are often strongly unfavorable energetically, particularly in condensed phases such as liquids, leading to a high rejection rate.

Two efficient methods that have been developed for particle insertion and deletion are the Unbonding–Bonding (UB) method^49,50^ and the AVBMC method.^23,24,36^ These two methods are nearly equivalent in efficiency and follow very similar schemes for particle insertion. Their deletion procedures differ slightly, with AVBMC offering a more symmetric formulation between insertion and deletion. In our simulations, we have employed AVBMC, although either method would have been equally suitable. In the AVBMC method, a molecule within the simulation box is first randomly selected as the target molecule. A new molecule is then inserted at a random position within the bound region—a spherical volume surrounding the target molecule. The insertion position can optionally be selected using the Rosenbluth scheme, weighted by the Boltzmann factor, to favor energetically favorable configurations. Thus, the AVBMC move accounts for both entropic contributions (through the volume of the bound region) and energetic contributions (via the Boltzmann weight).

For deletion, a target molecule is again randomly selected from the system. Then, one of its neighbors—i.e., a molecule located within its bound region—is randomly chosen and removed.

In cluster simulations, it is intuitively evident that particle insertion or deletion is more likely to occur at the cluster surface. Molecules near the surface tend to have fewer neighbors and, consequently, higher energies. Therefore, to enhance the efficiency of the AVBMC move, it is advantageous to replace uniform selection probabilities for (i) choosing the target molecule for insertion, (ii) choosing the target molecule for deletion, and (iii) choosing a neighbor of the target for deletion, with biased probabilities that favor molecules with higher energies.^25^

1. The probability of selecting molecule *i* as the target for insertion is given by:

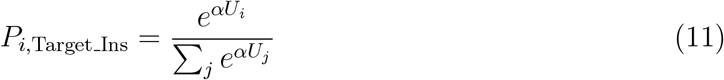

where *U*_*i*_ is the total interaction energy of molecule *i* with all other molecules, and *α* is a user-defined constant. Choosing *α* = *β* may overly bias the selection and exclude viable candidates. A moderate value, such as *α* = 0.15*β*, often provides a good balance between bias and diversity.
2. The probability of selecting molecule *i* as the target for deletion is given by:

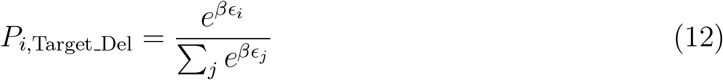

where *ϵ*_*i*_ is the energy of the highest-energy neighbor of molecule *i*.
3. Probability of selecting neighbor *i*^′^ of the target *i* for deletion:

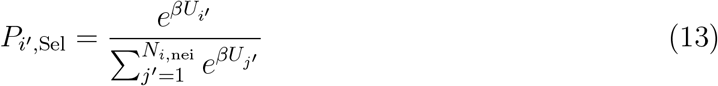

where *N*_*i*,nei_ is the number of neighbors of molecule *i*.

In our simulations of biomolecular clusters using the minimum-distance cluster criterion, insertion is performed by randomly selecting a segment of the inserting molecule and placing it within the bound region of a randomly chosen segment of the target molecule. The remainder of the molecule is then regrown segment by segment, with each step proceeding randomly toward either the N-terminal or C-terminal.

For deletion, the molecule to be removed is a neighbor of the target molecule. There are *N*_pair_ pairs of neighbored segments, each consisting of one segment from the target molecule and one from the neighboring (deleting) molecule. One of these pairs is randomly selected to initiate regrowth of the deleting molecule in order to calculate the generating probability for the old state.

The regrowth of a molecule in the dilute phase—which occurs during insertion (for evaluating the Rosenbluth weight of the old state) and during deletion (for evaluating the Rosenbluth weight of the new state)—is performed considering only intramolecular non-bonded interactions.

The acceptance probability for an insertion move, where the cluster size changes from *n* in the old state to *n* + 1 in the new state, is given by:

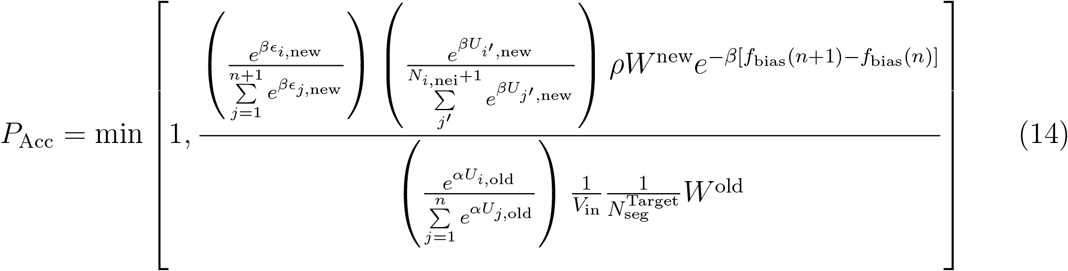

Here, *ρ* is the dilute-phase density, estimated by the ideal gas formula under the assumption that molecules do not interact in the dilute phase:

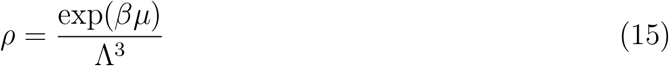

where 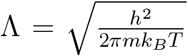 is the thermal de Broglie wavelength, *h* is Planck’s constant, and *m* is the molecular mass. *V*_in_ is the volume of the bound region, and 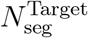 is the number of segments in the target molecule. The number of segments in the inserting molecule, 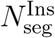, appears in both the old and new state terms and thus cancels out.

The acceptance probability for a deletion move, where the cluster size changes from *n* in the old state to *n* − 1 in the new state, is:

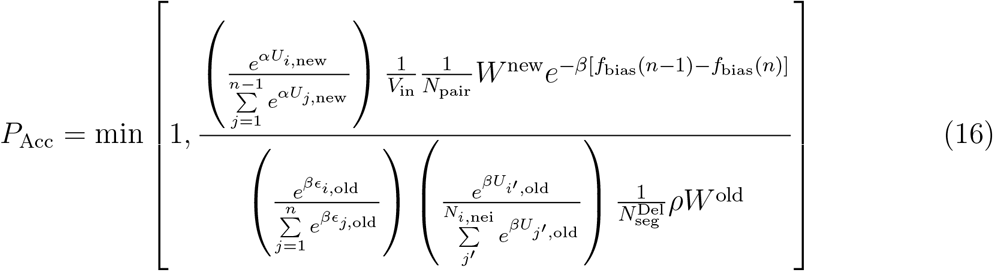

where 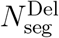 is the number of segments in the deleting molecule.

As stated in Eq. 3, the biasing potential for each cluster size *n* is updated after each MC iteration, where *N* (*n*) represents the cluster size histogram generated using 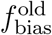. Ideally, the biasing potential converges after a few iterations, resulting in a uniform histogram across all cluster sizes. This has been successfully achieved in simulations of small molecules such as alkanes (C5–C7)^37^ and water.^38^ However, in our simulations involving biomolecules, producing a uniform histogram is significantly more challenging. This difficulty arises from the large size and flexibility of biomolecules, which often consist of many segments. In an insertion or deletion move, the entire molecule must be grown, leading to a very low acceptance probability. Consequently, in each MC iteration, some cluster sizes are sampled more frequently than others, resulting in fluctuations in the histogram.

One solution to this problem is to use the histogram data from all MC iterations to calculate the free energy of nucleation in a smoother and statistically consistent way. For this purpose, we employ the Weighted Histogram Analysis Method (WHAM). ^51^ WHAM combines the histograms collected under different biasing potentials and provides an optimal estimate of the unbiased probability distribution *P* (*n*). In its classical form, WHAM estimates *P* (*n*) as:

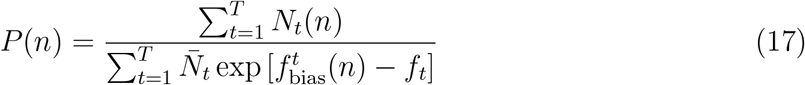

where *N*_*t*_(*n*) is the histogram count at cluster size *n* during iteration 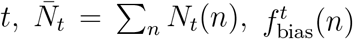 is the biasing potential at iteration *t*, and *f*_*t*_ is the free energy offset for that iteration.

The *f*_*t*_ values are solved self-consistently using:

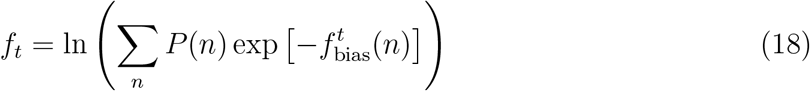

To reduce statistical noise from low-count bins and stabilize convergence, we use a modified version of WHAM, where the numerator is regularized as:

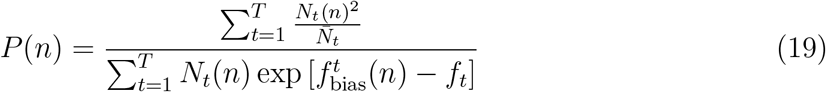

Once the unbiased probability *P* (*n*) is obtained, the free energy of nucleation is calculated as:

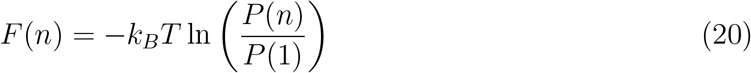

##### Force Fields

We applied our GCMC method by using two different coarse-grained force fields: multiscale *π* − *π* (MPIPI) model^41^ and a simplified hydrophobic-polar sidechain (HPS) model. ^17^ Both employ implicit solvent representations and treat each amino acid as a single interaction site (i.e., bead or segment), resulting in a linear chain model of the biomolecule. The only bonded interaction is bond stretching, modeled using a harmonic potential with fully flexible bond angles and torsions:

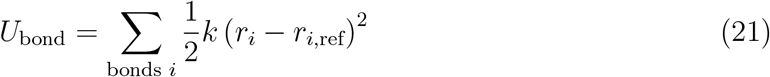

The spring constant *k* is 8.03 and 1.0 J mol^−1^ pm^−2^ for MPIPI and HPS, respectively. The reference bond length *r*_*i*,ref_ is 381 and 380 pm for MPIPI and HPS, respectively.

Non-bonded interactions, which apply to all pairs of non-bonded segments, include electrostatic and short-range potentials. In both force fields, electrostatic interactions between segments *i* and *j* separated by distance *r*_*ij*_ are modeled using a screened Coulomb potential based on the Debye–Hückel approximation:

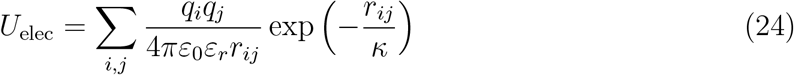

where *q*_*i*_ and *q*_*j*_ are the electrostatic charges of segments *i* and *j, ε*_0_ is the vacuum permittivity, *ε*_*r*_ = 80 is the relative dielectric constant, and *κ* is the Debye screening length. The assigned charges and electrostatic cutoffs differ between the two models: 3.5 nm for MPIPI and 4.0 nm for HPS.

In MPIPI, short-range pair interactions are modeled using the Wang–Frenkel (WF) potential:^52^

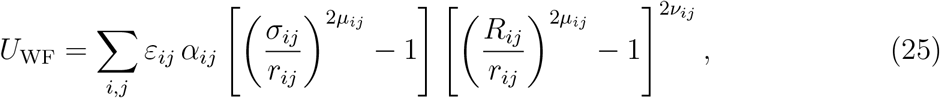

where *R*_*ij*_ = 3*σ*_*ij*_ is the interaction cutoff. The prefactor *α*_*ij*_ ensures that the potential smoothly decays to zero at the cutoff distance and is given by:

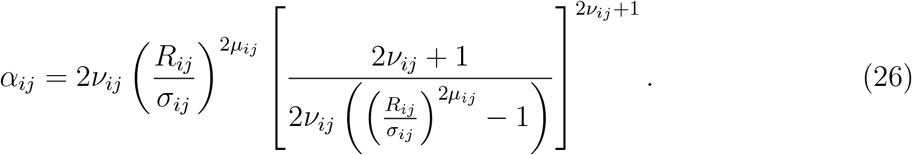

For the HPS model, a standard Lennard-Jones-like potential^17^ is used:

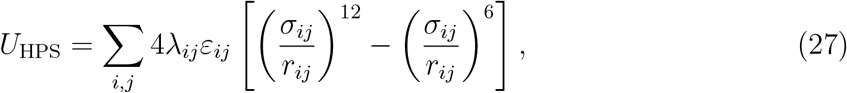

where *λ*_*ij*_ is a tunable hydrophobicity scaling factor. The cutoff for HPS interactions is set to 4.0 nm. Parameters for both force fields are provided in their respective references.

During chain growth in CBMC, FECBMC, or AVBMC moves, each trial for segment’s orientation is sampled in spherical coordinates by generating a bond length *r*, polar angle *θ*, and azimuthal angle *ϕ* from a distribution proportional to 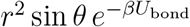.

The bond length is sampled from a Gaussian distribution with mean *r*_ref_ and standard deviation (*kβ*)^−1*/*2^ using the Box–Muller method,^53^ followed by an accept–reject step to ensure the correct *r*^2^ weighting.

In CBMC and AVBMC moves, the polar angle *θ*, defined as the angle between the new segment and the previous two segments, is drawn from sin *θ* over the interval (0, *π*), and the azimuthal angle *ϕ*, which defines the rotation about the bond axis, is drawn uniformly from (0, 2*π*).

In FECBMC moves, the polar angle *θ* is constrained to the interval (0, *θ*^max^), where *θ*^max^ ensures that the growing chain can geometrically connect to the second fixed point *F*. The Jacobian factor for growing *N*_*G*_ segments between two fixed points is:

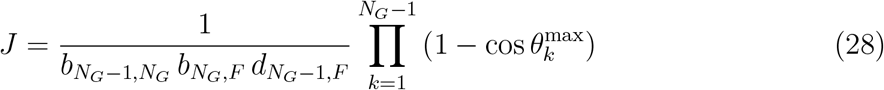

where 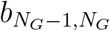 is the bond length between the last two grown segments, 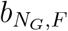 is the bond length between the last grown segment and the fixed endpoint *F*, and 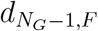 is the Euclidean distance between them. The azimuthal angle *ϕ*, which defines the rotation of the newly grown segment around the axis formed by the previous segment and the endpoint *F*, is sampled uniformly from the interval (0, 2*π*).

## Results and Discussions

We present two sets of results in this section. First, we benchmark the accuracy of our MC conformational sampling framework against established molecular dynamics (MD) data for single disordered proteins in dilute solution. This validation step ensures that the physical behavior of individual chains is faithfully captured in nucleation simulations. Second, we apply our GCMC scheme to map the nucleation free energy landscapes of two well-characterized intrinsically disordered proteins (IDPs), FUS-LC and NDDX4, using both HPS and MPIPI force fields. Our results reveal protein-specific differences in nucleation pathways, including a striking example of nonclassical behavior in NDDX4 under certain conditions.

We adapted the nucleation code developed by Troy Loeffler;^54^ incorporated the MPIPI and HPS force fields, a minimum atomic distance criterion for cluster identification, and additional features such as conformation sampling and swap moves tailored for biomolecular condensates.

### Comparison of MC and MD Simulations for Validation of Single Molecule Conformations

As an initial benchmark, we performed simulations on 17 proteins previously used to validate the MPIPI force field.^41^ These proteins span a wide range of amino acid compositions and electrostatic profiles, making them suitable test cases for assessing conformational sampling. The sequences of these proteins include various fractions of aromatic, positively charged, negatively charged, and neutral residues (Supporting Text)—with and without *π*-electron interactions. Joseph et al. conducted MD simulations on single protein molecules at *T* = 300 K, using Debye screening lengths adjusted to match available experimental data.^41^ They observed good agreement between experimental measurements and their computed radius of gyration (*R*_*g*_).^41^ The proteins and their corresponding (inverse) Debye screening lengths are listed in Table 1.

**Table 1:** Name, length (size), inverse Debye screening length (*κ*^−1^), and scaling exponents *ν* of proteins selected for MC and MD comparison.

| **No.** | **Name** | **Size** | $\kappa^{-1}$ ($\text{\AA}^{-1}$) | $\nu_{MD}$ | $\nu_{MC}$ |
| --- | --- | --- | --- | --- | --- |
| 1 | ACTR | 71 | 0.145 | 0.60 | 0.61 |
| 2 | α-Synuclein | 140 | 0.140 | 0.60 | 0.58 |
| 3 | hNHE1cdt | 131 | 0.145 | 0.58 | 0.59 |
| 4 | IBB | 97 | 0.131 | 0.55 | 0.55 |
| 5 | N49 | 36 | 0.131 | 0.61 | 0.61 |
| 6 | N98 | 151 | 0.131 | 0.55 | 0.56 |
| 7 | NLS | 44 | 0.131 | 0.62 | 0.62 |
| 8 | NSP | 176 | 0.131 | 0.54 | 0.54 |
| 9 | NUL | 112 | 0.131 | 0.56 | 0.57 |
| 10 | NUS | 80 | 0.131 | 0.57 | 0.58 |
| 11 | P53 | 93 | 0.149 | 0.59 | 0.60 |
| 12 | SH4-UD | 85 | 0.152 | 0.57 | 0.58 |
| 13 | Sic | 92 | 0.131 | 0.59 | 0.60 |
| 14 | Ash1 | 83 | 0.126 | 0.60 | 0.60 |
| 15 | K18 | 130 | 0.131 | 0.60 | 0.59 |
| 16 | K25 | 185 | 0.131 | 0.59 | 0.57 |
| 17 | ProTα | 111 | 0.128 | 0.65 | 0.66 |

We also located these proteins on Uversky^55^ and Das–Pappu diagrams,^56^ as shown in Figure 1. The Uversky diagram plots hydropathy, measured as the average Kyte–Doolittle score,^57^ against the mean net charge, defined as |*f*_+_ − *f*_−_|, where *f*_+_ and *f*_−_ represent the fractions of positively and negatively charged residues, respectively. The dashed line in the Uversky diagram approximately distinguishes globular proteins (above the line) from intrinsically disordered proteins (below the line).

**Figure 1:**
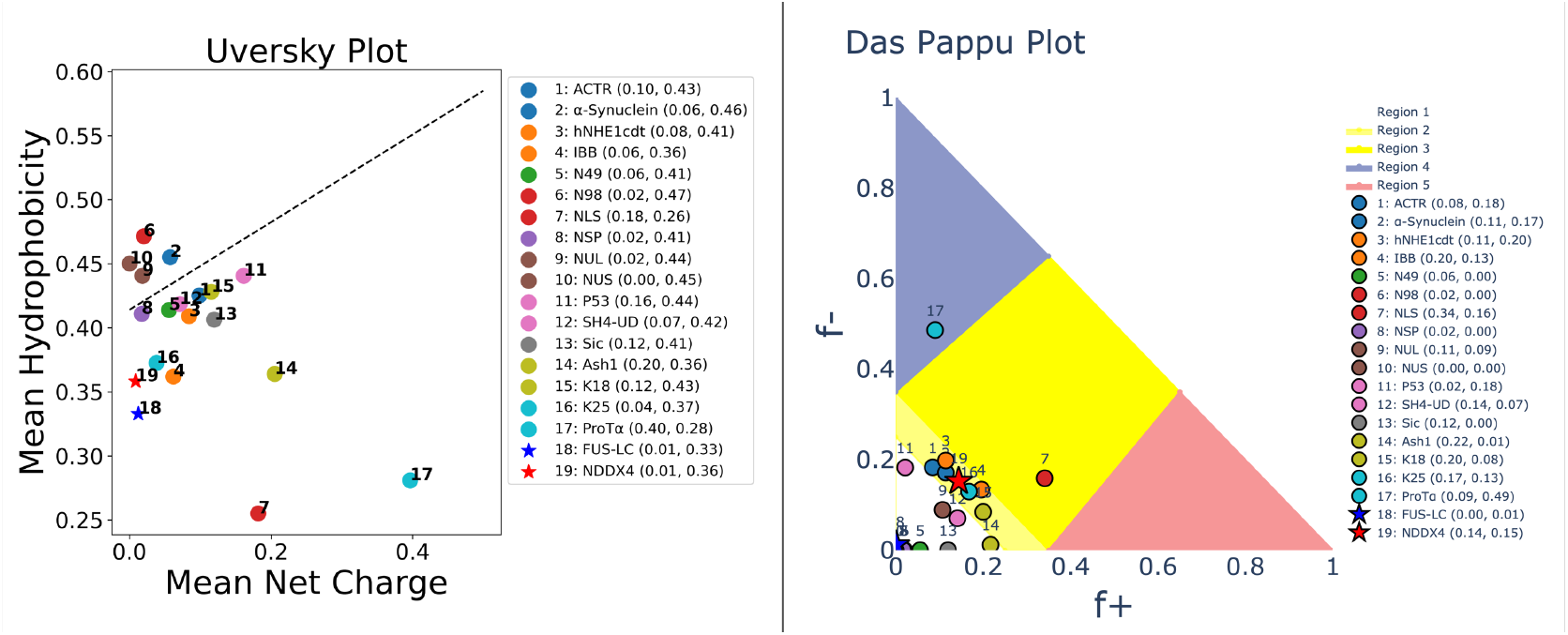
Uversky (left) and Das–Pappu (right) diagrams for 17 proteins selected for the single molecule simulations (as the numbers given in Table 1) (circles); and FUS-LC (18) and NDDX4 (19) (stars).

The Das–Pappu diagram classifies proteins based on *f*_+_ and *f*_−_ into five distinct regions: Region 1 corresponds to weak polyampholytes, Region 3 to strong polyampholytes, and Region 2 marks the boundary between these two. Regions 4 and 5 represent strong polyelectrolytes, proteins with highly negative and highly positive net charges, respectively. Our analysis confirms that the 17 proteins span diverse regions of both diagrams.

We repeated their MD simulations and also performed MC simulations for single molecules of each protein at *T* = 300 K using their respective Debye screening lengths. We only applied CBMC and FECBMC trial moves for the single-molecule MC simulations. Figures 2 and 3 compare the simulation results for the distribution of *R*_*g*_ and the ensemble-averaged intrachain scaling as a function of chain length (|*i* − *j*|).

**Figure 2:**
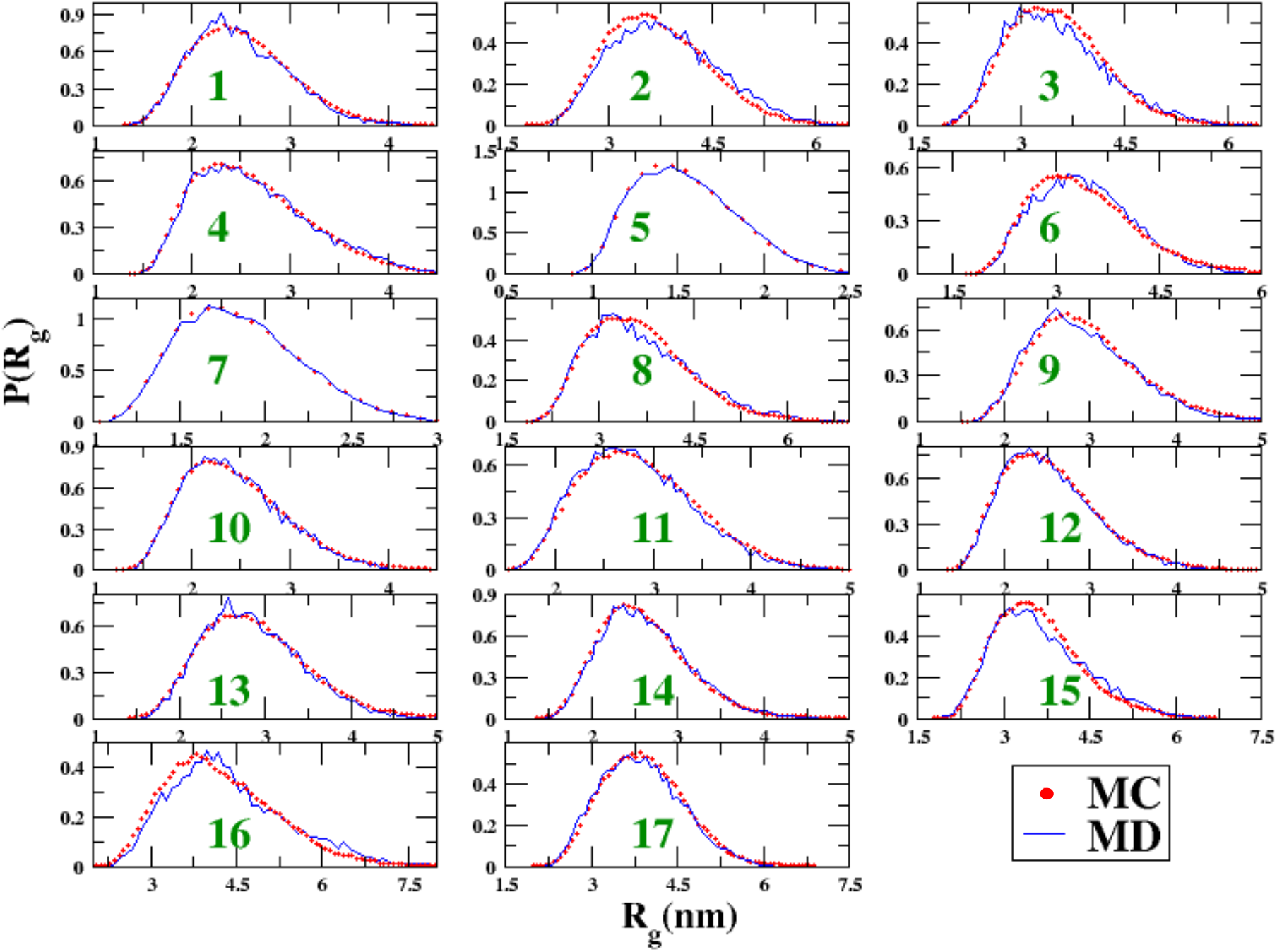
*R*_*g*_ distributions from MC and MD simulations for proteins 1 to 17 as labeled in Table 1.

**Figure 3:**
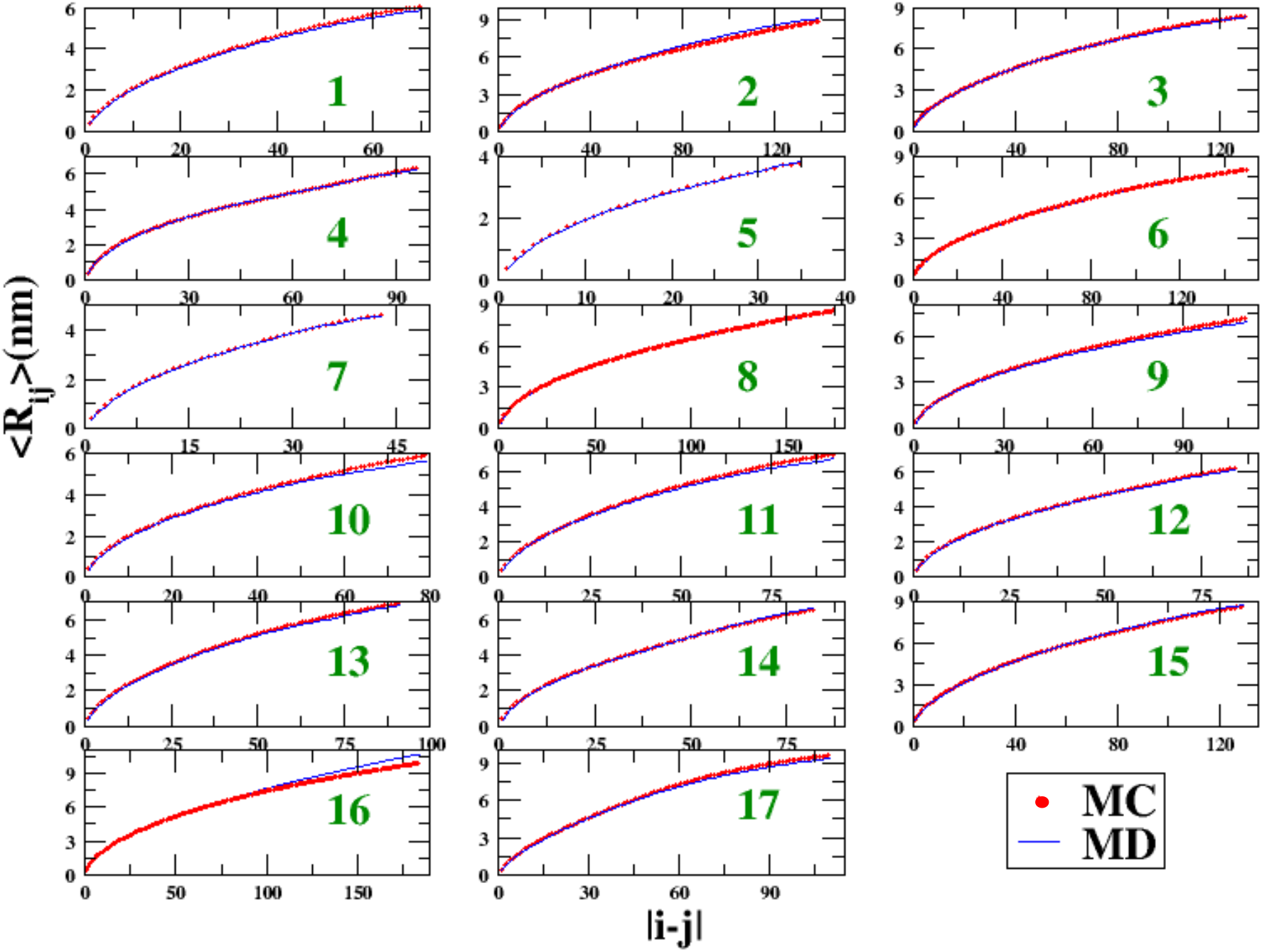
Intrachain distance between residues *i* and *j*, denoted ⟨*R*_*ij*_⟩, scaling as a function of chain separation |*i* − *j*| from MC and MD simulations for proteins 1 to 17 as labeled in Table 1.

Figure 2 shows that for all 17 proteins, the *R*_*g*_ distributions obtained from MC and MD simulations are in excellent agreement. We fitted the ensemble-averaged intrachain distances ⟨*R*_*ij*_⟩, to a power law function of sequence separation |*i* − *j*| as ⟨*R*_*ij*_⟩ = *A*|*i* − *j*|^*ν*^, where the exponent *ν* characterizes the solvent quality based on chain compactness:^58^ *ν* ≈ 0.6 indicates good solvation, *ν* ≈ 0.5 corresponds to theta conditions, and *ν* ≈ 0.33 represents poor solvation. The fitted exponents from both MC and MD simulations are listed in Table 1. The ⟨*R*_*ij*_⟩ profiles in Figure 3 and the exponent values in Table 1 demonstrate good consistency between the MC and MD results.

Since the exponents for all 17 proteins are in the proximity to the good solvation regime (*ν* for all *>* 0.5), we further examined two synthetic polyampholytes from Das and Pappu’s work,^56^ sv1 and sv30, at *T* = 300 K and inverse Debye screening length, *κ*^−1^ = 0. 126 Å^−1^. Each sequence contains 25 lysine (K) and 25 glutamate (E) residues, with sv1 arranged in an alternating sequence, (EK)_25_, and sv30 in a blocky pattern, E_25_K_25_. From the original work of Das and Pappu,^56^ *R*_*g*_ of sv1 is expected to be approximately 1 nm larger than that of sv30. The ⟨*R*_*ij*_⟩ profile for sv1 should increase monotonically, scaling with an exponent of 0.6. In contrast, sv30 is expected to adopt a compact hairpin-like conformation, resulting in a ⟨*R*_*ij*_⟩ profile that exhibits significant deviations from a simple power-law scaling behavior. The MC results in Figure 4 are consistent with these expectations and agree well with the results of the original work.^56^

**Figure 4:**
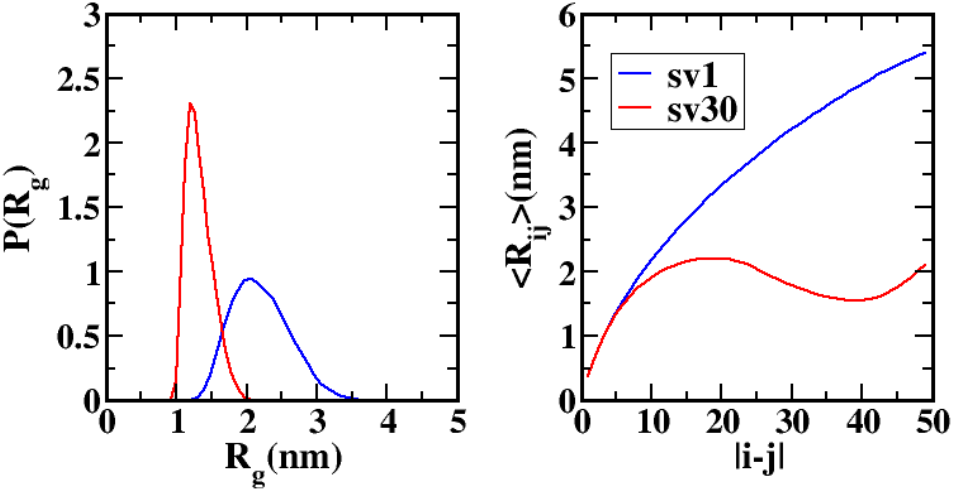
*R*_*g*_ distributions and ⟨*R*_*ij*_⟩ scaling as a function of chain separation |*i* − *j*| from MC simulations for polyampholytes sv1 and sv30.

**Figure 5:**
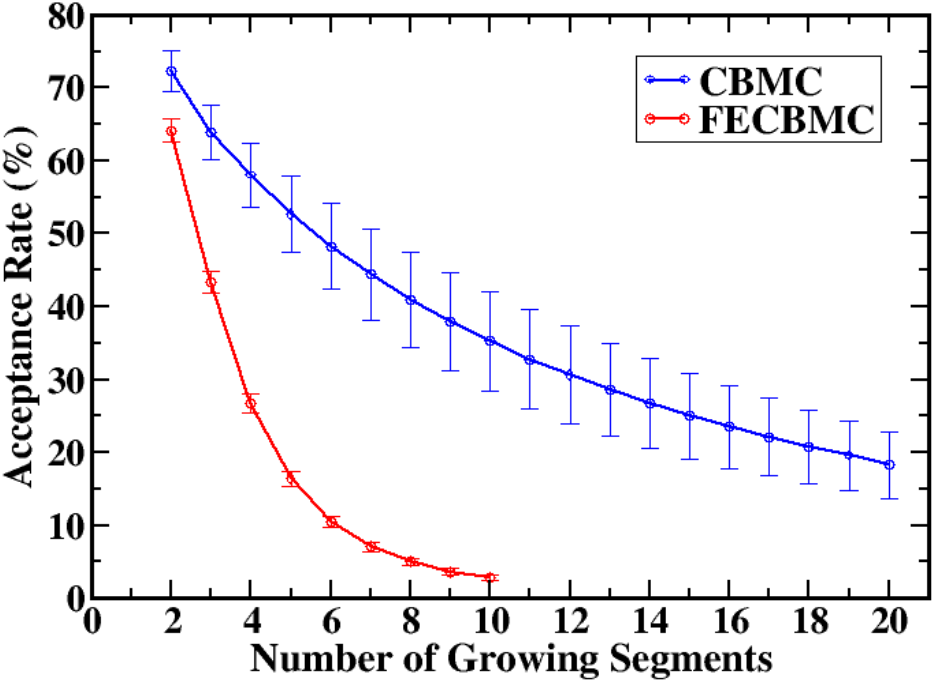
Acceptance rates of CBMC and FECBMC moves as a function of the number of growing segments.

In our MC simulations of a single molecule, the number of trial orientations generated during CBMC and FECBMC moves was set to 10, which is computationally inexpensive. Figure 5 shows the acceptance rates of CBMC and FECBMC moves as functions of the number of growing segments for 10 trial moves. For both CBMC and FECBMC moves, the acceptance rate decreases as the number of growing segments increases. This trend is expected, as the overall acceptance probability is the product of the individual acceptance probabilities for each growing segment (see Eq. 8).

Moreover, the acceptance rate of FECBMC is consistently lower than that of CBMC for the same number of growing segments and declines more rapidly. This difference arises because CBMC imposes no constraints on the final position of the growing chain, allowing greater flexibility to explore low-energy conformations. In contrast, FECBMC requires the chain to be closed between two fixed points, which restricts conformational sampling and reduces the likelihood of finding energetically favorable structures.

Overall, the data presented in Figure 5 demonstrate that both CBMC and FECBMC are efficient and computationally viable methods for conformational sampling. Given that these results are for large proteins with extensive intramolecular nonbonded interactions, it is expected that the acceptance rates for sampling conformations of individual molecules within clusters will not be significantly lower.

The cumulative results presented in this section—from Table 1 to Figures 1 through 5—demonstrate that our MC framework can accurately and efficiently sample the conformational landscapes of a diverse range of proteins.

### FUS-LC and NDDX4 Nucleation Free Energy Surfaces

NDDX4 and FUS-LC are well-established examples of intrinsically disordered proteins (IDPs) that undergo LLPS to form biomolecular condensates involved in diverse aspects of cellular organization. Experimental studies have shown that the phase behavior of these proteins is governed by a complex interplay of sequence-encoded interactions, including cation–*π* and *π*–*π* contacts, modulated by sequence charge patterning, aromatic content, temperature, and salt concentration.^6,59^ FUS-LC, comprising 163 residues and a molecular weight of 17,149.75 g/mol, is enriched in aromatic (24 Tyr) and polar (42 Ser and 37 Gln) residues, which promote *π*–*π* stacking and polar interactions that drive its phase separation. In contrast, NDDX4 contains 236 residues with a molecular weight of 25,412.48 g/mol and exhibits a high density of charged residues organized in alternating blocks of positively charged (24 Arg and 8 Lys) and negatively charged (18 Glu and 18 Asp) segments, making it a strong polyampholyte. Our analysis of the sequence on the Uversky diagram (Figure 1, left) confirms that both proteins are located on the lower hydrophobicity side, consistent with the presentation of IDPs. The analysis on the Das-Pappu diagram (Figure 1, right) shows that while FUS-LC lies in the weak polyampholyte region, NDDX4 is close to the boundary of strong polyampholytes.

Previous MD simulations have investigated the nucleation behavior of these proteins using a modified liquid droplet model and HPS model.^17^ Here, we apply our GCMC framework to simulate the nucleation of NDDX4 and FUS-LC under identical conditions: *T* = 300 K and *κ* = 10 Å.

We performed our MC simulations using both MPIPI and HPS force fields for NDDX4 and FUS-LC to investigate the effect of force field on the nucleation free energy profile. The maximum cluster size is set to *n*_max_ = 100, and the dilute density *ρ* is chosen to be sufficiently low to ensure that the critical nucleus size exceeds 100.

In each MC step, one move is selected according to the following probabilities: conformation sampling (0.5), translation (0.23), rotation (0.23), and swap (0.04). Although the final converged result is independent of these probabilities, their careful tuning can significantly accelerate convergence.

When a conformation sampling move is chosen, a random molecule is selected, and a section of the chain is randomly chosen for segment rearrangement. If this section lies at one end of the chain, a CBMC move is used; otherwise, an FECBMC move is performed. When a swap move is selected, insertion or deletion is attempted with equal probability. In such cases, the entire molecule must be regrown using CBMC moves for both the old and new states. As shown in Figure 5, the acceptance rate of CBMC moves decreases with increasing the number of growing segments. Therefore, many trial orientations (e.g., 100– 500) is required to achieve a satisfactory acceptance rate for swap moves. Longer chains require more trial orientations, and simulations with the MPIPI force field typically demand more trials than those with HPS.

Each MC cycle consists of 100 MC moves, providing every molecule in the cluster a reasonable chance to be updated. To ensure adequate sampling of all cluster sizes, each MC iteration must contain a sufficiently large number of cycles, typically between 100,000 and 200,000.

Figure 6 (a)-(d) show the nucleation free energy profiles of NDDX4 and FUS-LC obtained from GCMC simulations using both the HPS and MPIPI force fields at lower supersaturation.

**Figure 6:**
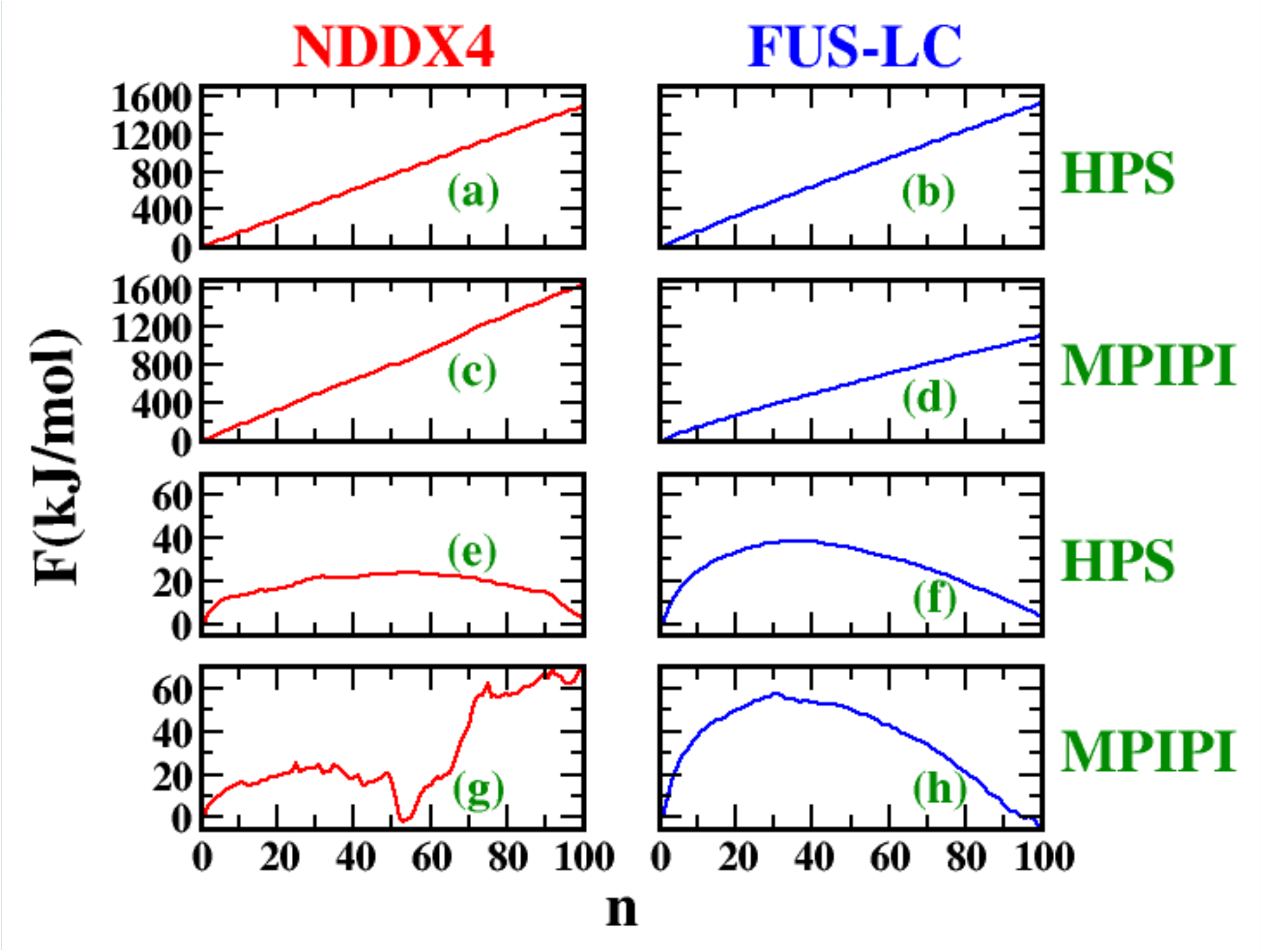
Free energy profiles for (a) NDDX4 with HPS force field at 0.844 g/cm^3^, (b) FUS-LC with HPS force field at 0.57 g/cm^3^, (c) NDDX4 with MPIPI force field at 0.105 g/cm^3^, (d) FUS-LC with MPIPI force field at 1.424 g/cm^3^, (e) NDDX4 with the HPS force field at 337.6 g/cm^3^, (f) FUS-LC with the HPS force field at 262.2 g/cm^3^, (g) NDDX4 with the MPIPI force field at 63 g/cm^3^, and (h) FUS-LC with the MPIPI force field at 128.16 g/cm^3^.

Each of these free energy profiles, obtained at a low dilute density, can be transformed to correspond to any arbitrary dilute density using Equation 4. For our analysis, we selected dilute densities that yield a free energy barrier at cluster sizes below 100, as illustrated in Figure 6 (e)-(h).

According to Figure 6 (e)-(h), for equal free energy barriers, both NDDX4 and FUS-LC require lower dilute densities when simulated with the MPIPI force field compared to the HPS force field. While both force fields exhibit a classical nucleation profile for FUS-LC—characterized by a single free energy maximum—the nucleation behavior differs for NDDX4. The HPS force field yields a classical nucleation profile, whereas the MPIPI force field indicates a nonclassical nucleation pathway, with a metastable intermediate state appearing near a cluster size of ∼53. Indeed, Figure 6 (g) reveals a single free energy barrier up to cluster size ∼53, consistent with classical nucleation. However, beyond cluster size ∼53, a second barrier emerges, indicating a departure from classical behavior.

To better understand the structural origins of the nonclassical nucleation behavior observed in NDDX4, we analyzed representative cluster configurations along the nucleation coordinate (Figure 7). For FUS-LC, cluster morphology evolves smoothly with size and remains roughly spherical throughout, consistent with the assumptions of classical nucleation theory. In contrast, NDDX4 (with MPIPI force field) exhibits qualitatively distinct behavior. Up to the metastable state at cluster size 53, clusters remain compact and spherical. However, beyond this size, configurations increasingly resemble two partially fused subclusters, indicating a failure to coalesce into a single, isotropic nucleus.

**Figure 7:**
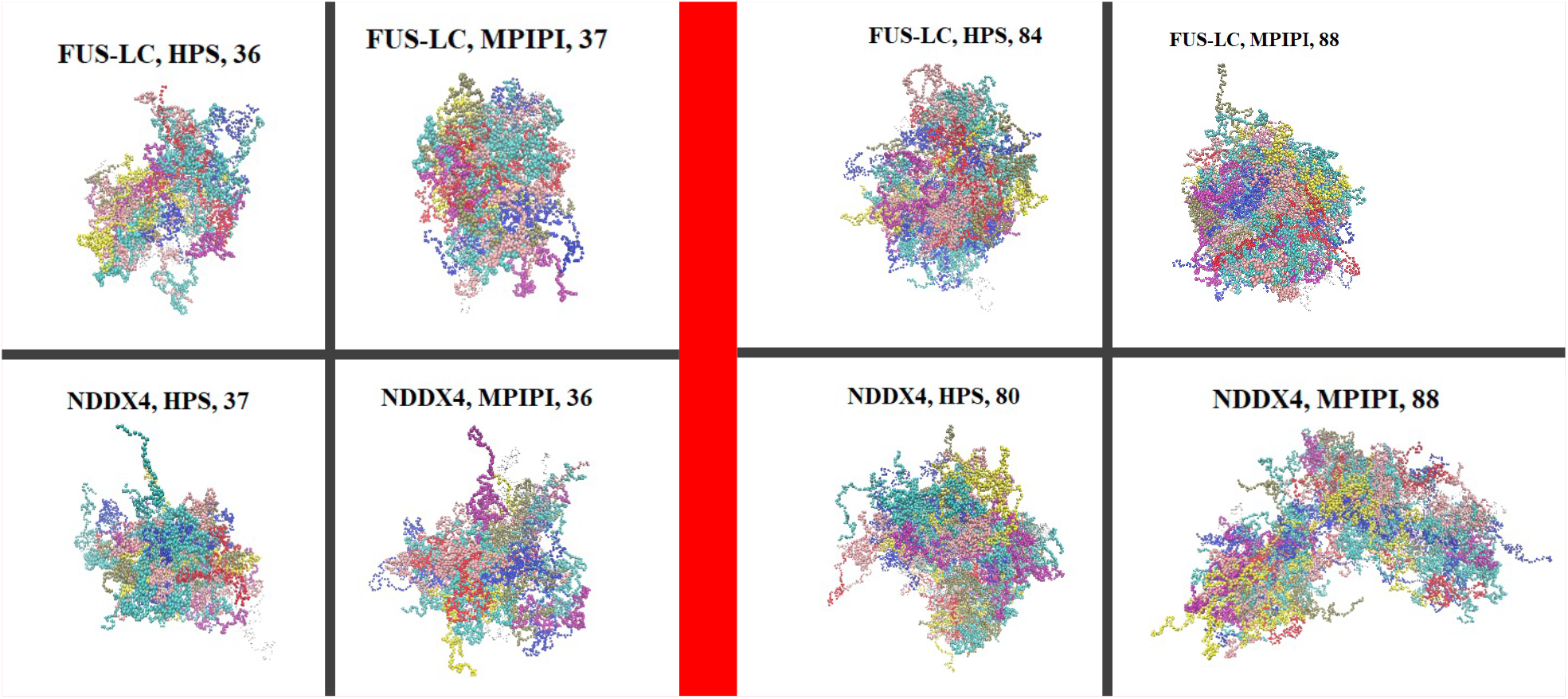
Representative snapshots of molecular clusters from GCMC simulations at small (∼36, left panel) and large (∼85, right panel) cluster sizes for FUS-LC and NDDX4 using HPS and MPIPI force fields. Molecules are rendered in distinct colors to aid visual differentiation, although a limited palette necessitated color repetition. The snapshots illustrate classical growth for all systems except NDDX4 with MPIPI, where the ∼85-molecule structure appears as two loosely connected subclusters, consistent with a metastable intermediate and a nonclassical nucleation mechanism.

This morphological transition suggests that the nucleation barrier in NDDX4 is not purely thermodynamic, but also involves geometric frustration—possibly related to electrostatic repulsion between charged segments or difficulties in reorganizing domain-level interactions. These multi-lobed structures likely represent metastable intermediates stabilized by sequence-specific interactions. Such features are fundamentally beyond the scope of classical nucleation models, which assume spherical nuclei with smooth growth. Their spontaneous emergence in our GCMC simulations highlights the strength of our approach: it does not rely on predefined assumptions and can reveal complex nucleation pathways driven by molecular detail.

Previous studies^41^ have shown that the MPIPI force field more accurately captures phase separation phenomena than HPS-type force fields—for example, in reproducing the phase behavior of poly-Arg and poly-Lys peptides in the presence of RNA. In our simulations, MPIPI also promotes the formation of stable NDDX4 clusters, likely driven by electrostatic and cation–*π* interactions that are largely absent in FUS-LC.

Compared to previous MD simulations^17^ (performed only with the HPS force field), our GCMC results are consistent in predicting that, at comparable dilute densities, FUS-LC exhibits a higher nucleation barrier than NDDX4. However, our MC simulations yield significantly larger free energy barriers. This discrepancy arises primarily from the different cluster definitions used in the two approaches. In our simulations, a minimum atomic distance criterion with a 5 Å cutoff is employed—a method previously used in conjunction with Martini force fields.^45^ Further adaptation of this criterion to our one-bead-per-residue model is warranted and will be guided by radial distribution function analysis in future work. In contrast, the MD study assumes the presence of a single large cluster representing the condensed phase, with the remaining proteins forming the dilute phase. A two-dimensional clustering criterion—based on interchain distances and the distance of each protein’s center of mass from the condensed phase—is used in that approach.

## Conclusion

In this work, we developed an MC simulation framework to simulate the nucleation of biomolecular condensates and map their nucleation free energy surfaces. A key advantage of our approach is the use of the grand canonical ensemble, which allows the number of particles in the system to fluctuate throughout the simulation. This feature eliminates the finite-size effect observed in canonical ensemble simulations, where the dilute-phase concentration decreases as molecules accumulate in clusters.

A central strength of our method is its ability to independently sample each cluster size and resolve both classical and nonclassical nucleation pathways without imposing prior assumptions. We demonstrated that our conformational sampling moves—based on CBMC and FECBMC algorithms—can efficiently explore the conformational landscape of single molecules, yielding accurate estimates for properties such as the radius of gyration and ensemble-averaged intrachain distance across a diverse set of proteins.

We further applied this framework to study the nucleation behavior of two representative intrinsically disordered proteins, NDDX4 and FUS-LC, using both MPIPI and HPS force fields. In simulations involving swap moves—where an entire molecule must be regrown—many trial orientations (e.g., 100–500) are necessary to achieve satisfactory acceptance rates (1–5%), highlighting the need for continued development of more efficient growth strategies.

When applied to FUS-LC and NDDX4, two phase-separating IDPs, our method uncovered force field–dependent differences in nucleation mechanisms. While both proteins exhibited classical nucleation behavior with the HPS force field, only FUS-LC remained classical with MPIPI. NDDX4, in contrast, showed a nonclassical nucleation landscape under MPIPI, featuring a metastable intermediate near cluster sizes 53. Morphological analysis revealed that clusters beyond this point adopt nonspherical, multi-lobed structures—indicative of frustrated coalescence and barriers to spherical fusion. This behavior underscores the intricate coupling between molecular sequence, interaction specificity, and nucleation dynamics.

Our findings highlight the critical importance of force field selection and sequence design in determining nucleation pathways. More broadly, they demonstrate that direct sampling of nucleation landscapes—rather than fitting to classical models—can reveal previously inaccessible mechanistic detail. Looking ahead, a major strength of the grand canonical ensemble is its natural extensibility to multicomponent systems, making it a promising platform for simulating binary^60^ and ternary^61^ nucleation. In future work, we plan to extend this framework to study the competitive nucleation between multiple protein species, the role of co-factors such as RNA, and the influence of sequence patterning in heterogeneous mixtures. These extensions will enable deeper insight into the sequence-dependent rules that govern phase behavior and nucleation pathways in complex biological environments.

## Acknowledgement

This work was supported by funding from the Cancer Prevention and Research Institute of Texas (CPRIT) award RR220008, the Welch Foundation (Award E-2221 and Catalyst Center for Advanced Bioactive Materials Crystallization Award V-E-0001), and NSF CBET-2442006. The simulations presented in this work were performed using the computational resources provided by the Hewlett-Packard Enterprise Data Science Institute at the University of Houston.

## Supporting Information Available

Sequences of all selected proteins are provided in the Supporting Information.

## TOC Graphic

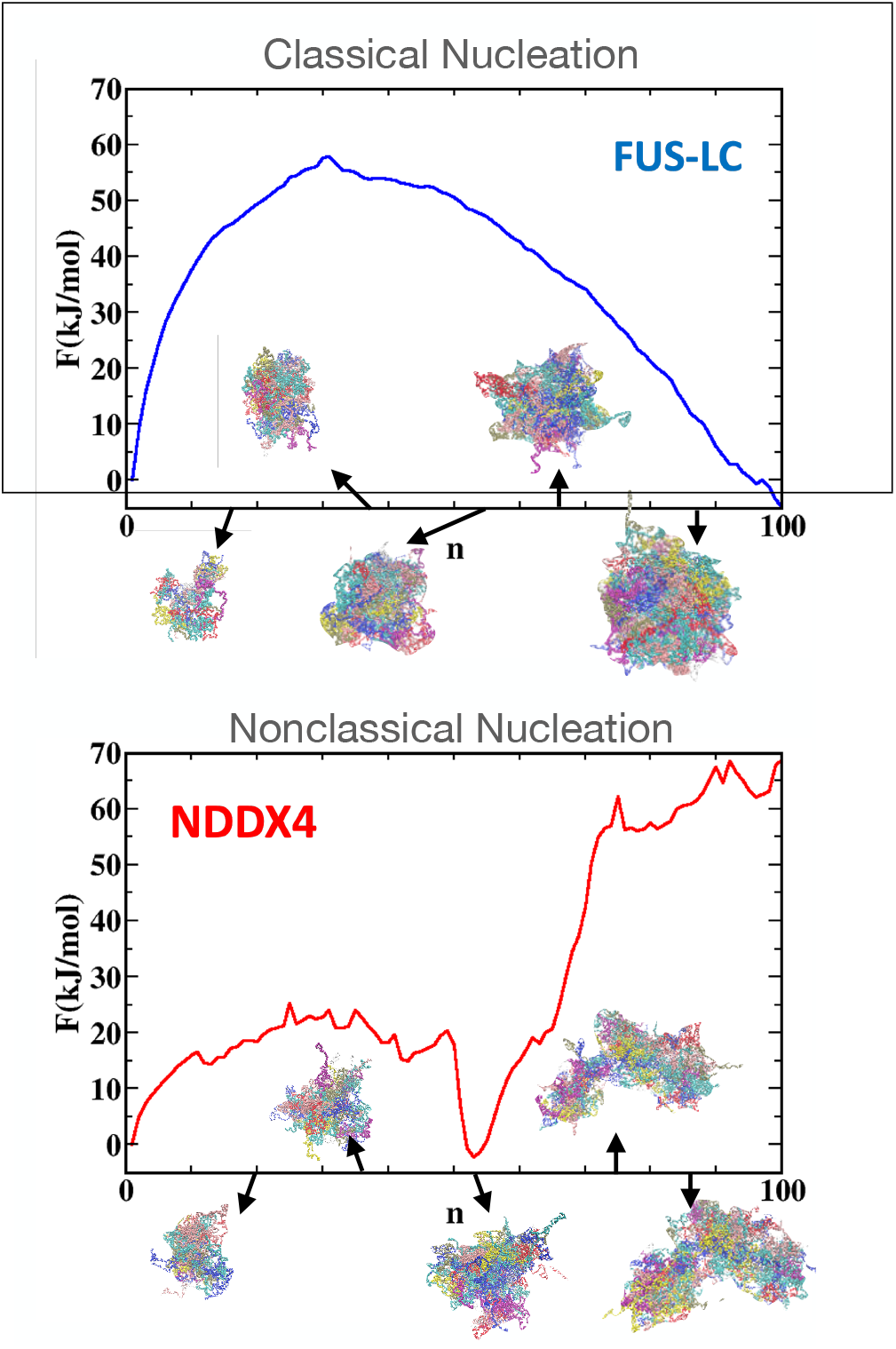

