## Supporting Information for "Nucleation Landscape of Biomolecular Condensates in the Grand Canonical Ensemble via Monte Carlo Simulations"

### Supporting Text

#### Sequences of the selected proteins

**ACTR:**

GTQNRPLLRNSLDDLVGPPSNLEGQSDERALLDQLHTLLSNTDATGLEEIDRALGIPELVN QGQALEPKQD

**$\alpha$-Synuclein:**

MDVFMKGLSKAKEGVVAAAEKTKQGVAEAAGKTKEGVLYVGSKTKEGVVHGVATVAE KTKEQVTNVGGAVVTGVTAVAQKTVEGAGSIAAATGFVKKDQLGKNEEGAPQEGILEDM-PVDPDNEAYEMPSEEGYQDYEPEA

**hNHE1cdt:**

MVPAHKLDSPTMSRARIGSDPLAYEPKEDLPVITIDPASPQSPESVDLVNEELKGKVLG

LSRDPAKVAEEDEDDDGGIMMRSKETSSPGTDDVFTPAPSDSPSSQRIQRCLSDPGPH-
PEPGEGEPFFPKGQ

**IBB:**

GCTNENANTPAARLHRFKNKGKDSTEMRRRRIEVNVELRKAKKDDQMLKRRNVSSFPDD
ATSPLQENRNNQGTVNWSVDDIVKGINSSNVENQLQAT

**N49:**

GCQTSRGLFGNNNTNNINNSSSGMNNASAGLFGSKP

**N98:**

GCFNKSFGTPFGGGTGGFGTTSTFGQNTGFGTTSGGAFGTSAFGSSNNTGGLFGNSQTKP
GGLFGTSSFSQPATSTSTGFGFGTSTGTANTLFGTASTGTSLFSSQNNAFAQNKPTGFGN-
FGTSTSSGGLFGTTNTTSNPFGSTSGSLFGP

**NLS:**

ACETNKRKREQISTDNEAKMQIQEEKSPKKKRKKRSSKANKPPE

**NSP:**

GCNFNTPQQNKTPFSFGTANNNSNTTNQNSSTGAGAFGTGQSTFGFNNSAPNNTNNANSS
ITPAFGSNNTGNTAFGNSNPTSNVFGSNNSTTNTFGSNSAGTSLFGSSSAQQTKSNGTAG-
GNTFGSSSLFNNSTNSNTTKPAFGGLNFGGGNNTTPSSTGNANTSNNLFGATANAN

**NUL:**

GCGFKGFDTSSSSSNSAASSSFKFGVSSSSSGPSQTLTSTGNFKFGDQGGFKIGVSSDSGSIN
PMSEGFKFSKPIGDFKFGVSSESKPEEVKKDSKNDNFKFGLSSGLSNPV

**NUS:**

GCPSASPAFGANQTPTFGQSQGASQPNPPGFGSISSSTALFPTGSQPAPPTFGTVSSSSQ
PPVFGQQPSQSAFGSGTTPN

**P53:**

MEEPQSDPSVEPPLSQETFSDLWKLLPENNVLSPLPSQAMDDLMLSPDDIEQWFTEDPGP
DEAPRMPEAAPPVAPAPAAPTPAAPAPAPSWPL

**SH4-UD:**

MGSNKSKPKDASQRRRSLEPAENVHGAGGGAFPASQTPSKPASADGHRGPSAAFAPAAA
EPKLFGGFNSSDTVTSPQRAGPLAGG

**Sic:**

GSMTPSTPPRSRGTRYLAQPSGNTSSSALMQGQKTPQKPSQNLVPVTPSTTKSFKNAPLLA
PPNSNMGMTSPFNGLTSPQRSPFPKSSVKRT

**Ash1:**

GASASSSPSPSTPTKSGKMRSRSSSPVRPKAYTPSPRSPNYHRFALDSPPQSPRRSSNSSIT
KKGSRRSSGSSPTRHTTRVCV

**K18:**

MQTAPVPMPDLKNVKSKIGSTENLKHQPGGGKVQIINKKLDLSNVQSKCGSKDNIKHVPGGG
SVQIVYKPVDLSKVTSKCGSLGNIHHKPGGGQVEVKSEKLDFKDRVQSKIGSLDNITHVPGGG
NKKIE

**K25:**

MAEPRQEFEVMEDHAGTYGLGDRKDQGGYTMHQDQEGDTDAGLKAEEAGIGDTPSLEDEA
AGHVTQARMVSKSKDGTGSDDKKAKGADGKTKIATPRGAAPPGQKGQANATRIPAKTP
PAPKTPPSSGEPPKSGDRSGYSSPGSPGTPGSRSRTPSLPTPPTREPKKVAVVRTPPKSPSS
AKSRL

**ProT$\alpha$:**

MSDAAVDTSSEITTKDLKEKKEVVEEAENGRDAPANGNAENEENGEQEADNEVDEEEEE
GGEEEEEEEEGDGEEEDGDEDEEAESATGKRAAEDDEDDDVDTKKQKTDEDD

**sv1:**

EKEKEKEKEKEKEKEKEKEKEKEKEKEKEKEKEKEKEKEKEKEKEKEKEK

**sv30:**

EEEEEEEEEEEEEEEEEEEEEEEEEKKKKKKKKKKKKKKKKKKKKKKKKK

**NDDX4:**

MGDEDWEAEINPHMSSYVPIFEKDRYSGENGDNFNRTPASSSEMDDGPSRRDHFMKSGFA
SGRNFGNRDAGECNKRDNTSTMGGFGVGKSFGNRGFSNSRFEDGDSSGFWRESSND-

CEDNPTRNRGFSKRGGYRDGNNSEASGPYRRGGRGSFRGCRGGFGLGSPNNDLDPDECMQ

RTGGLFGSRRPVLSGTGNGDTSQSRSGSGSERGGYKGLNEEVITGSGKNSWKSEAEGGES

**FUS-LC:**

MASNDYTQQATQSYGAYPTQPGQGYSQQSSQPYGQQSYSGYSQSTDTSGYGQSSYSSYGQ

SQNTGYGTQSTPQGYGSTGGYGSSQSSQSSYGQQSSYPGYGQQPAPSSTSGSYGSSSQSSSYGQ

PQSGSYSQQPSYGGQQQSYGQQQSYNPPQGYGQQNQYNS
